# OmniNA: A foundation model for nucleotide sequences

**DOI:** 10.1101/2024.01.14.575543

**Authors:** Xilin Shen, Xiangchun Li

**Author notes:** Correspondence to: Prof. Xiangchun Li, Tianjin Cancer Institute, Tianjin Medical University Cancer Institute, and Hospital, Huanhu Xi Road, Tiyuan Bei, Hexi District, Tianjin, 300060, China.

## Abstract

Foundation models have demonstrated exceptional efficacy across diverse downstream tasks. However, within the realms of genomics and transcriptomics, a notable gap persists in the availability of models that afford a comprehensive understanding of nucleotide sequence principles across various species. Here, we present OmniNA, a foundation generative model designed for comprehensive nucleotide sequence learning. The model was pre-trained on 91.7 million nucleotide sequences and the corresponding annotations encompassing 1076.2 billion bases and 197 million words spanning a multitude of species. We demonstrated OmniNA gains the capacity to understand the semantics of the nucleotide sequence and textual annotations by analyzing the learned representation of the pre-trained model. OmniNA can be fine-tuned to align multiple nucleotide learning tasks with natural language paradigms. We demonstrate OmniNA-1.7B surpasses or rivals state-of-the art methods in 17 nucleotide tasks, encompassing nucleotide sequences detection and species classification. The model’s understanding of nucleotide grammars enhances its capability to reveal the mutation effect of nucleotide sequence on DNA and RNA processing. We hereby release the OmniNA-1.7B model as an open-source contribution to the research community. This foundation model signifies a step toward advancing our comprehension of nucleotide sequences across diverse species and holds substantial promise to facilitating genomics and transcriptomics research.

## Introduction

Artificial intelligence (AI) is undergoing a paradigm shift to the foundation model that leverages large language model (LLM) incorporating millions of parameters trained on extensive data, versatile for a wide range of predictive applications ^1^. These models learn the underlying data patterns and structures from massive unlabeled data, which facilitate innovative applications via transfer learning ^2–4^. It has emerged as a transformative technology with many successful examples in various fields, such as the Generative Pre-trained Transformer (GPT) ^5^ in natural language processing and Stable Diffusion ^6^ in vision language. In the ever-evolving landscape of genomics, the advent of the foundation model has emerged as a transformative force, propelling our capacity to decode the intricate language embedded within nucleic acid sequences. Against this backdrop, we present a large-scale generative pre-trained foundation model tailored specifically for nucleic acid sequences across diverse species and biological contexts. This initiative harnesses the power of deep learning in genomics, where the burgeoning complexity of biological data demands innovative solutions for deciphering the underlying grammar governing nucleotide sequences.

The rapid progress of deep learning methodologies in genomics has been propelled by the escalating volume and diversity of genomic data. Existing models, such as Enformer and TIGER ^7,8^, have made contributions to specific genome tasks. Enformer leverage a transformer architecture ^9^ for expression and chromatin states prediction. TIGER is a deep convolutional network to predict off-target activity of RNA-targeting CRISPR guide RNA sequence ^8^. However, these deep learning approaches are tailored to specific genome tasks, and thus have limited generalization ability to other tasks. Additionally, as the model parameters are trained from scratch with labeled data, they cannot be transferred across tasks. In the ever-evolving landscape of genomic data, it is crucial to develop foundation models capable of adapting to various nucleotide sequences, subsequently transferring the knowledge to a broad spectrum of genomic tasks.

This study proposes OmniNA, a foundation model for nucleotide sequence. OmniNA was pre-trained on a scale of 91.7 million (M) nucleotide sequences encompassing 1076.2 billion (B) bases range across a global species and biological context. Unlike previous works, our approach integrates nucleotide sequence and the corresponding text annotations into the learning process. To our knowledge, this is the first study to learn so far the largest publicly available nucleotide sequences spanning a diverse array of species. By doing so, the model gains the capacity to understand semantic of text annotations, thereby empowering it to perform a wide range of downstream tasks. Our analysis demonstrates the model learned the syntax of the nucleotide sequence by analyzing the learned representations of sequences. Next, a comprehensive evaluation of our pre-trained model was conducted to assess its ability to adapt to new captions to handle multi-tasks in one model. After fine-tuning, the model gains capability to unveil the mutational effects from its comprehension of nucleotide grammars.

## Methods

### Data processing for OmniNA pre-training

We curate sequence data along with corresponding textual annotations from the Nucleotide (nt) Database of the National Center for Biotechnology Information (NCBI) up to April, 2023. These data encompass genomic DNA and RNA sequences derived across diverse species. Employing a sliding window approach, we truncate sequences longer than 3000 bp to 3000 bp. Sequences with length less than 200 bp are excluded. We filter out sequences containing equal to or more than two consecutive N nucleotides. Subsequently, we obtain a total of 91, 732, 311 sequences, collectively comprising 1, 076, 183, 408, 570 bases. The textual annotations have a total of 197, 089, 040 words. We design the prompts to organize the sequence data with the text annotations as feature-ordered input for model pre-training (**Supplementary Table 1**). The prompt is composed of three modules: 1. an instruction; 2. a nucleotide sequence; 3. an annotation.

### Model architecture and pre-training

We established OmniNA model with the LLaMA architecture ^10^, which is an auto-regressive decoder-only transformer adopted the self-attention mechanism. The data were tokenized with the byte pair encoding (BPE) algorithm ^11^. We train the tokenizer using nucleotide sequences and texts of wiki-103 dataset ^12^. SentencePiece ^13^ is employed as the subword tokenizer, resulting in a dictionary containing 32, 001 tokens. Subsequently, the nucleotide sequences and text annotations are partitioned into sentence-based chunks. We include a special token <S= at the beginning and a special token at the end of the sequence. Each chunk is further tokenized using the trained tokenizer. The max length of input sequence is set to 601 tokens. The token embedding layer embedded the input tokens into embedding vectors. We add GPTNeo positional encodings ^14^ to the token embeddings. The transformer consists of multiple layers, each containing a multi-head self-attention layer and a feedforward layer. We construct three models with varying heads and layers, which range from 66 M up to 1.7 B parameters (**Supplementary** Figure 1). The transformer layer captures token-token interaction information using multi-head self-attention mechanism^9^. Given an input, the self-attention mechanism assigns an attention weight α*_i_*_,*j*_ > 0 to each token pair *i*,*j*, where ∑*_j_* α*_i_*_,*j*_ = 1. Attention in the transformer is bidirectional. Attention weights α*_i,j_* are computed by the scaled dot-product of the query vector of *i*(*Q*) and the key vector of *j*(*K*), followed by a softmax operation. Then attention weights are used to produce a weighted sum of value vectors (*V*):

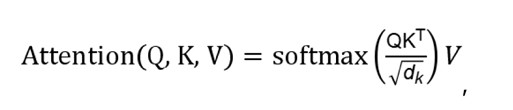

where *d_k_* is the dimension of *K*. Before feeding into transformer layers, we normalize the input with root mean square layer normalization ^15^. We replace the ReLU non-linearity of the traditional transformer architecture by SwiGLU activation function ^16^. The output of the last transformer layer is fed into a linear layer for auto-regressive learning ^5^. The sequence and annotations were trained in an auto-regression. Specifically, the prediction at each time step depends on the preceding observations. For example, the prediction for the base *b_t_* at time *t* is based on the preceding observation *b_t-1_* at time *t*−1:

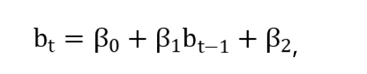

where β*_0_*, β*_1_* and β*_2_* are trainable parameters governing the auto-regressive prediction. Mean Squared Error (MSE) loss function was applied for auto-regression learning, defined as:

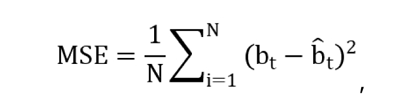

where *N* represents the number of tokens in one input. A detailed model architecture is shown in **Supplementary** Figure 2.

The model is trained using the AdamW optimizer ^17^, with the following hyper-parameters: β*_1_* = 0.9, β*_2_* = 0.999. We use a learning rate of 1e-4, and a batch size of 2, 048 for 200, 000 steps (**Supplementary** Figure 1). The learning rate is decayed with a cosine schedule, and a warm-up ratio of 0.03. We use a weight decay of 0.0001 and gradient clipping of 1.0. All experiments were run in python v.3.8. Detailed software versions are: PyTorch v.1.12.1; CUDA v.11.7; CUDNN v.8.5.0; transformer v.4.28, datasets v.2.11 and tokenizers v.0.13. The model was run on an NVIDIA DGX A100 system with 8 GPUs, each with 80GB of memory.

### Evaluation of OmniNA’s generation ability

We primarily focus on the fidelity, diversity, randomicity and stability of the generations of the pre-trained OmniNA. The genomic contents are generated with beam search if unspecifically. We first evaluate the model fidelity as the consistency between OmniNA generated annotation and the ground truth annotation. Three annotation tasks, including species, genomic types and enzyme annotations, are applied for the evaluation. We consider exact matches as correct after applying a vocabulary mapping that converts model-predicted vocab to informal names and abbreviations. We collected 172 species that appeared in the NCBI nt database for more than 1, 000 times. We randomly sampled 100 sequences for each of the 172 species. For genome type annotation task, 100 sequences for each of the coding sequence, intron, genome coding region, mRNA, snRNA, lncRNA, miRNA, rRNA, ncRNA, and tRNA are randomly sampled from the NCBI nt database. The informal names and abbreviations of the taxonomies are collected from the GenBank taxonomy database (https://www.ncbi.nlm.nih.gov/taxonomy). For enzyme annotation task, we collected 104 enzymes that appeared in the NCBI nt database for more than 1, 000 times. We randomly sampled 100 sequences for each of the 104 enzymes from the NCBI nt database. We also evaluate the fidelity of the sequence generation as the similarity between the generated sequence and the ground truth sequence. The similarity was evaluated as the cosine similarity between the token embedding of the sequences. The bacteria and virus coding sequence are collected from the NCBI nt database.

We further quantified the conservation of protein coding sequences as the mean cosine similarity among the sequences’ token embeddings of the respective protein among different taxonomies. The cosine similarity ranges from -1 to 1, where higher values suggest greater similarity. The diversity of tokens was quantified with Shannon entropy from 10, 000 randomly sampled sequences from the NCBI nt database. We consider exact matches as correct after applying a vocabulary mapping that converts model-predicted vocab to informal names and abbreviations. The Shannon entropy score ranges from 0 to a maximum value, where higher values suggest greater diversity and unpredictability of the tokens in a given set. The official gene names are collected from the NCBI database (https://ftp.ncbi.nih.gov/gene/). The reproducibility of model-generated text annotation is quantified as the similarity among the different repeats of the model generations. We randomly select 10, 000 sequences from the NCBI nt library and generate annotations for 10 repeats using nucleus sampling generation algorithm ^18^ with a top_p = 0.92. The reproducibility is quantified as the mean cosine similarity among the token embeddings of the 10 repeats. We next evaluate the stability of the model generated annotations, which is assessed as the consistency of the generated annotations among different truncations of the same sequence. We randomly sampled 10, 000 sequences with lengths > 1, 000 from the NCBI nt database and randomly truncated them into 10 fragments per sequence. The stability is quantified as the mean cosine similarity among the feature representations of the annotations.

### Feature representation extraction from the pre-trained model

For each token presented to OmniNA-1.7B, the model embeds it into a 2048-dimensional vector that encodes the characteristics of the token to the sequence context. We extracted the embedding vector of the vocab and nucleotide sequence from the token embedding layers of the pre-trained OmniNA-1.7B model. For each token, the average embedding vectors of the sub-tokens for each vocab or sequence are visualized with t-SNE dimensionality reduction.

For each transformer layer, we further extract the feature representations of the nucleotide sequence as the output of transformer layers. The adjusted rand index (ARI) score is applied to evaluate the transformer layers in terms of sequence representation ability. The ARI score measures the percentage of matches between the sequence clusters and labels. The resulting score ranges from -1 to 1, where a higher score denotes that the data point fits well in the current cluster. We used the Louvain community detection algorithm implemented in “tl.louvain” of Scanpy package (version 1.7.0) for sequence clustering. In our study, Louvain algorithm would generate much more sequence clusters than real sequence types when the resolution was 1 and far fewer when the resolution was 0.0001. Thus, we set the resolution parameter range from 0.0001 to 1 with a step of 0.001 and computed ARI score with sklearn package (version 1.2.1) for each step. The maximum ARI score is employed as the final evaluation index. Three specific tasks in terms of sequence conservation delineation, species differentiation and functional sequence differentiation are applied for test. The sequence conservation of human genome is calculated with the GERP score ^19^. Specifically, employing a sliding window approach, we truncated the genomes into non-overlapped sequences of 1, 000 bp and calculated the mean GERP score for each sequence. Sequences with the highest and lowest quartile of GERP scores are defined as high and low conserved sequences, respectively. Bacteria and archaea 16s RNAs and fungi ITS regions were collected from the NCBI RefSeq database (https://www.ncbi.nlm.nih.gov/refseq/targetedloci/). The nucleotide sequences of SARS-Cov-2 were collected from GISAID (https://nextstrain.org/ncov/gisaid/global/6m). The coding sequence of functional proteins of bacteria and virus were extracted from the NCBI nt database. The promoter sequences are collected as the 2, 000 bp around the transcription start site of human GRCh38 reference genome. The human silencer and enhancer sequences are collected from the SilencerDB and VISITA enhancer browser ^20–22^. The 5’ and 3’ UTR sequences are collected from human GRCh38 reference genomes. The transcriptome factor binding sequences are randomly sampled from TF ChIP-seqs of the Encyclopedia of DNA Elements (ENCODE) database ^23^.

### Description of the downstream tasks

The model underwent fine-tuning for 17 tasks in a multi-task framework. **Supplementary Table 2** provides an overview of data resources and the splitting for training, validation, and test sets. Below, we briefly outline these tasks. We focused on tasks related to human DNA elements, encompassing promoter, enhancer, silencer, insulator, and active promoter and enhancer identification. Promoter sequences, defined as 2000 bp around the transcription start site were collected from GRCh38 and mm10 reference genomes. Enhancer, silencer and CTCF sequences were sourced from VISITA enhancer browser, SilencerDB and CTCFBSDB, respectively. Active promoter data were collected from MPRA-DragoNN. Active enhancer data were collected from DeepSTARR ^24^. The data for PAS signal, TIS signal, splice site and CRISPR off-target prediction were collected from previous studies ^8,25,26^. The genome-wide methylome profiles of 39 human cell types were sourced from a previous research ^27^. For chromatin accessibility, histone occupation and transcription factor (TF) binding site prediction, we downloaded the ATAC-seq, DNase-seq and ChIP-seq in .bed format from the ENCODE database (**Supplementary Table 3**). We retained data meeting two criteria: 1) data with the highest audit tiers defined in the ENCODE database or two biological replicates; 2) data with more than 1, 000 positive peaks. A total of 114 chromatin accessibility profiles from 89 cell lines were collected for chromatin accessibility prediction. A total of 884 TF ChIP-seq samples targeting 271 TFs across 56 cell lines were collected for TF binding prediction. A total of 248 histone ChIP-seq samples targeting 22 regions across 70 cell lines were collected for histone binding prediction. We defined a region with coverage RGB score <50 as a positive peak. We further filtered out peaks less than 200 bp or longer than 3000 bp. To balance data, we randomly sampled 6, 000 and 1, 000 peaks if the positive peak count overmuch for training and test, respectively. We further collected 193, 414 single nucleotide variants (SNVs) from the Clinvar database ^28^ that annotated with the “likely benign”, “benign”, “likely pathogenic” or “pathogenic”. For each SNV, we collected sequences length with 1, 000 bp around the mutation site for pathogenic or benign classification. Genes with more than 2 benign mutations and 2 pathogenic mutations were selected for the test. The data for the pathogenic virus prediction task are collected from DeePaC ^29^. The data for taxonomy prediction task are collected from DeepMicrobes ^30^.

### Apply the model for multi-task fine-tuning

**Supplementary Table 1** shows a prompt example for each task, which is composed of four modules: 1. an instruction; 2. a nucleotide sequence; 3. a test question; 4. an answer. The first two parts are fixed for all tasks, while the question and answer are task-specific. In the fine-tuning stage, only the answer was trained in an auto-regressive manner. The model is fine-tuned using the AdamW optimizer for 2 epochs ^17^, with the following hyper-parameters: β*_1_* = 0.9, β*_2_* = 0.999. The learning rate and batch size are varied with the size of the model. The learning rate is decayed with a cosine schedule, and a warmup ratio of 0.03.

### Benchmark OmniNA with task-specific deep learning methods

To assess OmniNA’s QA performance, we conducted a comparative evaluation against state-of-the-art deep learning methods across the 17 fine-tuning tasks with the test set (**Supplementary Table 4**). The benchmark methods employed for specific tasks are detailed in **Supplementary Table 4**. For the detection of human and mice DNA elements, including promoters, enhancers, silencers, and insulators, we employed benchmark methods such as PromID ^31^, iEnhancer-DEEP ^32^, DeepSilencer ^20^ and CTCFBSDB ^33^. Activation promoter and enhancer tasks were benchmarked with MPRA-DragoNN ^24^ and DeepSTARR ^34^. PAS signal, TIS signal and splice site detection tasks were benchmarked with DeepGSR ^25^ and SpliceRover ^35^. CRISPR-Cas13d off-target effects were benchmarked with TIGER ^8^. Chromatin accessibility, histone occupation, and TF binding prediction tasks were benchmarked with Enformer ^7^, Sei ^36^, and Expecto ^37^. Pathogenic SNP detection tasks were benchmarked with EVE ^38^. Pathogenic virus classification and species detection tasks were benchmarked with the DeePaC model ^29^. DeepMicrobes ^30^.

A total of 10 indexes are applied for the evaluation including precision, recall, F1 score, accuracy, false positive rate (FPR), true positive rate (TPR), false negative rate (FNR), true negative rate (TNR) and Matthew’s correlation coefficient (MCC). In multi-label classification tasks such as histone, TF and bacteria taxonomy classification, prediction accuracy for each label is individually assessed, treating any class with a different label as a negative sample.

### Saturation mutation analysis

We quantify mutation effects using log-likelihood (LL) differences between mutated (*s^mut^*) and wild-type (*s^WT^*) nucleotide sequences:

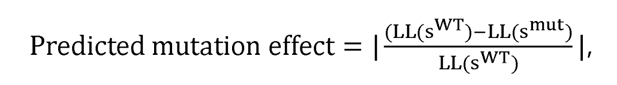

where the LLs are estimated using the OminHelix-1.7b model, representing the predicted LL of the positive target in the respective fine-tuning task:

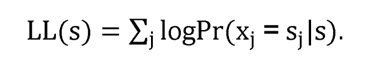

Benign and pathogenic mutations within the DNA sequences of PAS and TIS signal regions are sourced from the Clinvar database ^28^. The actual CRISPR-Cas13d off-target effect for each gRNA position is calculated as:

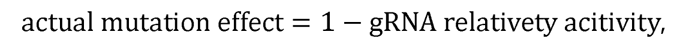

Where gRNA relativity activity is collected from a previous study ^8^. Domain regions of the BRCA1 and CHD7 proteins are obtained from the UniProt database ^8^. Full-length structures of BRCA1 and CHD7 spanning residues 200-1200 are predicted using AlphaFold2 ^39^.

### Statistical analyses

We generally employed t-test for the statistical analysis if unspecified. The paired t-test is used for assessing the differences between paired samples. Data distributions were determined through a permutation test involving 1, 000 permutations. For each permutation, the selected sample numbers were based on the data size. A significance threshold of *p*-value < 0.05 is adopted for all tests, and these are two-sided in nature. We adjust *p*-values using the False Discovery Rate (FDR) method to account for multiple comparisons.

## Data availability

The data for pre-training is openly available at https://ftp.ncbi.nlm.nih.gov/blast/db/FASTA/. The fine-tuning data are documented in **Supplementary Table 2**.

## Results

### An overview of OmniNA

We developed OmniNA as a foundation model for nucleotide sequence (**Figure 1**). The cognitive core of OmniNA is a LLM developed through incremental training on an extensive nucleotide sequence corpus, which inherits the benefits of emergent abilities of LLM and domain-specific knowledge. The backbone of OmniNA is a transformer-based auto-regressive decoder with self-attention mechanism ^10^. Three models with varying layer and head numbers were developed, spanning scales from 66 M to 330 M and up to 1.7 B parameters (**Supplementary** Figure 1). The detailed network structure is delineated in **Supplementary** Figure 2. We curated a dataset for model pre-training, which comprises 1076.2 B bases from 91.7 M nucleotide sequences across diverse species, extracted from the NCBI nt database (**Figure 1A**; see **Method**). We designed prompts to structure the sequences and their corresponding natural language descriptions as model input (**Figure 1A**; **Supplementary Table 1**). The input underwent chunking into subword tokens, followed by embedding through the token embedding layer. Token embeddings were then concatenated with position embeddings and fed into the transformer layers and followed by a regression layer for auto-regressive training (**Figure 1A and Supplementary** Figure 2) ^5^. The pre-trained OmniNA can handle versatile downstream applications in a multi-task manner (**Figure 1B**). After fine-tuning, the model can manage taxonomy and genomic element identification tasks in a Question-Answer (QA) paradigm (**Figure 1A**).

**Figure 1.**
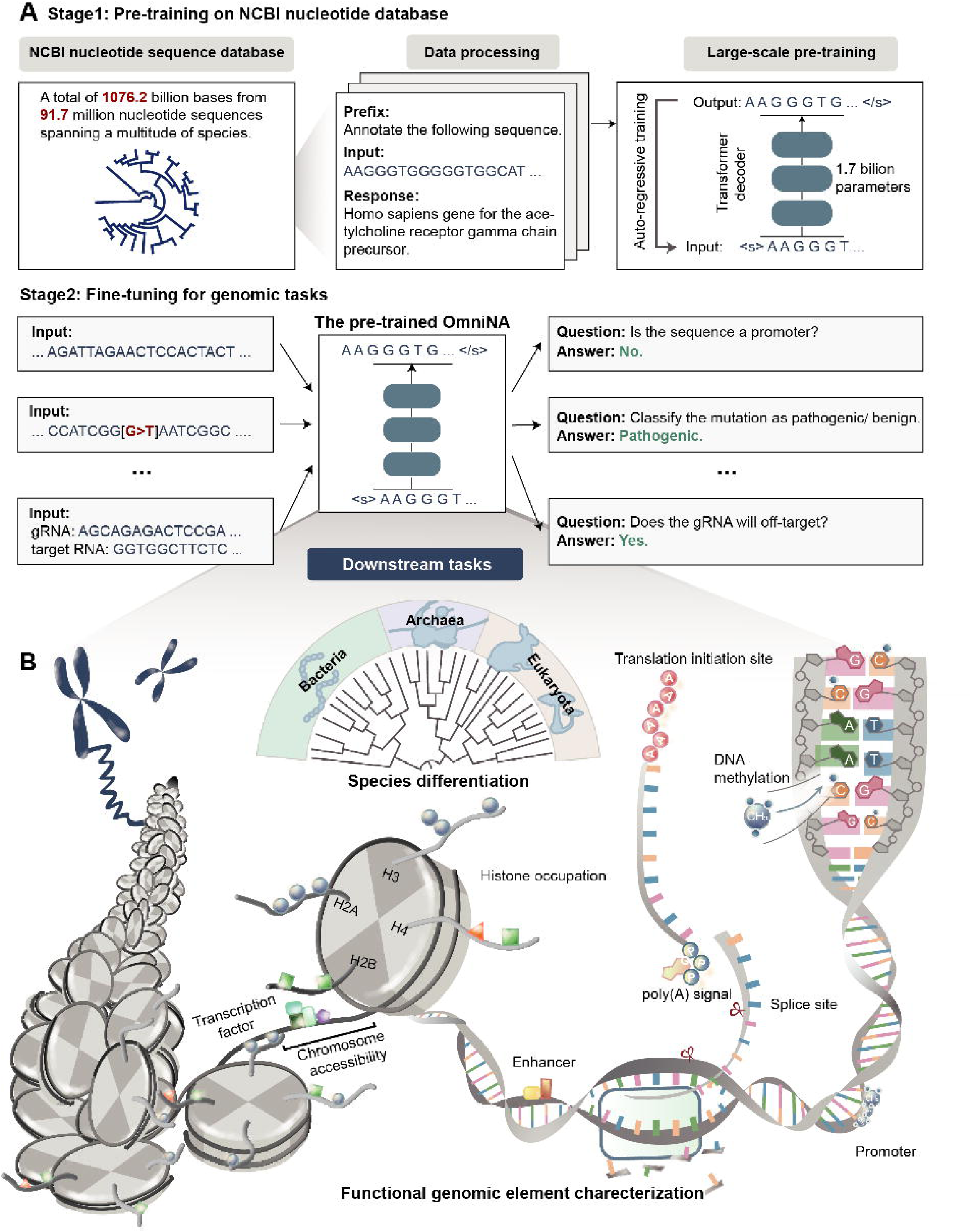
Schematic of OmniNA. (**A**) Nucleotide sequences, sourced from the NCBI nt database, are amalgamated with their corresponding text annotations for model pre-training. The model, featuring a transformer-based decoder, undergoes pre-training through an auto-regressive approach. Fine-tuning is executed across 17 tasks in a Question-Answer (QA) framework. (**B**) Overview of the downstream tasks encompassing nucleotide sequence detection, ranging from taxonomy classification to genomic element detection.

### The generation performance of OmniNA

In this section, we conducted a systematic evaluation of OmniNA’s generation capabilities across four aspects, including the fidelity, diversity, reproducibility and stability of the generated contents. The fidelity evaluation focused on the model’s ability to generate sequences and annotations consistent with ground truth counterparts. For sequences exhibiting high conservation (cosine similarity > 0.65), OmniNA consistently produced highly similar sequences across different scales. For sequences exhibiting low conservation (cosine similarity < 0.65), the largest model shows the most notable improvement in fidelity (**Supplementary** Figure 3). To further evluate fidelity in annotation generation, three annotation tasks were employed, including species, genomic type, and enzyme annotations. We observed an upward trend in generations’ F1 score with the increasing model size (**Figures 2A-B**), with OmniNA-1.7B achieving the highest F1 scores of 0.66, 0.24, 0.59 for species, genomic type and enzyme annotation tasks, respectively (**Figures 2A**, **Supplementary** Figure 3). Diving into diversity assessment, larger models demonstrated an increased capacity to generate diverse tokens, reflected in elevated token numbers and Shannon entropy (**Figures 2C-D**). Particularly, larger models exhibited a heightened likelihood of generating low-frequency words (**Figures 2C-D**). Our reproducibility analysis revealed that larger models exhibited higher consistency in generating annotations across different repeats, evident in increased cosine similarity (**Figure 2E**). Visual inspection indicated that sentences generated with OmniNA-1.7B with similar semantics were more likely to cluster together, and annotations for the same sequence were closer (**Figure 2F**). Evaluation of generation’s stability indicated that different truncations of the same sequence were more prone to yield similar annotations (**Figure 2G**). For instance, sequences from the same genomes of HIV produced more similar annotations than those from different genomes (**Figure 2H**). Notably, the model consistently generated correct annotations in both non-overlapping and overlapping regions of gag and pol genomes of HIV. In summary, our comprehensive evaluation reveals that the OmniNA model excels in consistently generating realistic and reasonable annotations.

**Figure 2.**
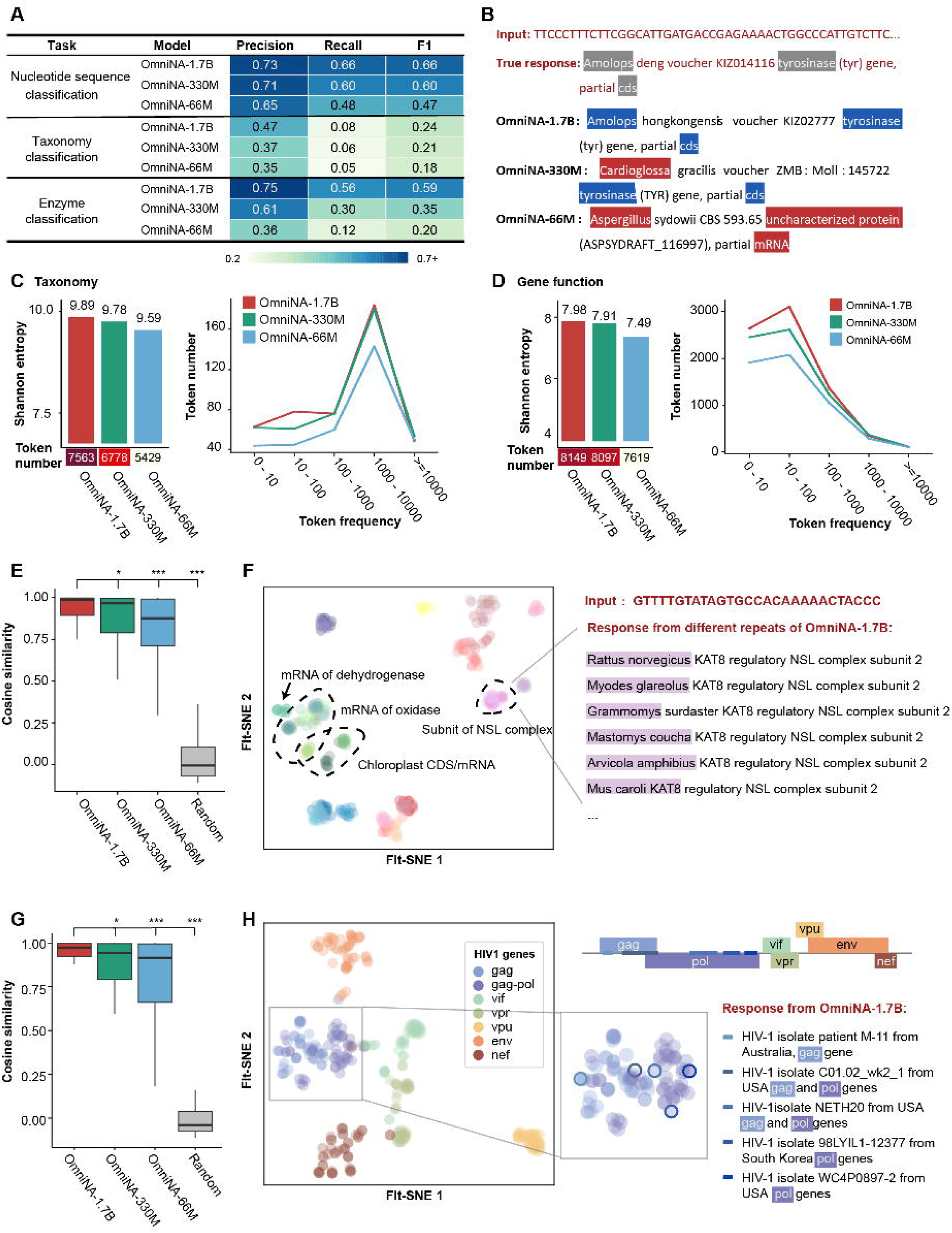
Evaluation of OmniNA’s generation performance. (**A**) Fidelity assessment in sequence annotations generated by OmniNA models. For each target and model, the precision, recall and F1 score represent the mean score across all labels. (**B**) Example of generated annotations for the same input by OmniNA models of varying sizes. (**C**) (Left) Bar plot illustrating Shannon entropy for taxonomy token generation. The heatmap at the bottom represents the generated token numbers. (Right) Line chart depicting generated token numbers correlated with token frequency. (**D**) (Left) Bar plot illustrating Shannon entropy for gene function token generation. The heatmap at the bottom represents the generated token numbers. (Right) Line chart depicting generated token numbers related to token frequency. (**E**) Box plot showcasing the cosine similarity of feature representations among different annotations generated from the same sentence. (**F**) t-SNE plot deciphering feature representations of generated annotations, with annotations from the same sequence colored identically. (**G**) Box plot reflecting the cosine similarity of token embeddings among different annotations generated from different truncations of the same sequence. (**H**) t-SNE plot deciphering the feature representations of generated annotations, with annotations from the same genome colored identically. \*\*\**p*-value < 0.001; \*\**p*-value < 0.01; \**p*-value < 0.05.

### Enhanced sequence and annotation representations by OmniNA

To gain a deeper understanding of OmniNA’s capabilities in aligning sequence and textual patterns with natural language, we investigated the token representations of the pre-trained OmniNA-1.7B (see **Methods**). t-SNE plot reveals the model’s ability to discriminate tokens representing distinct semantics, such as genus, genes and sequences (**Figure 3A**). Besides, tokens for gene symbols and gene functions exhibit distinct clustering, while sequence tokens with different base preferences manifest in separate loci (**Figure 3A**). Moreover, taxonomy names are distinctly separated, with tokens from the same taxonomy clustering together (**Figure 3B**). We also observed that viruses from different phyla form distinct subclusters (**Figure 3B**).

**Figure 3.**
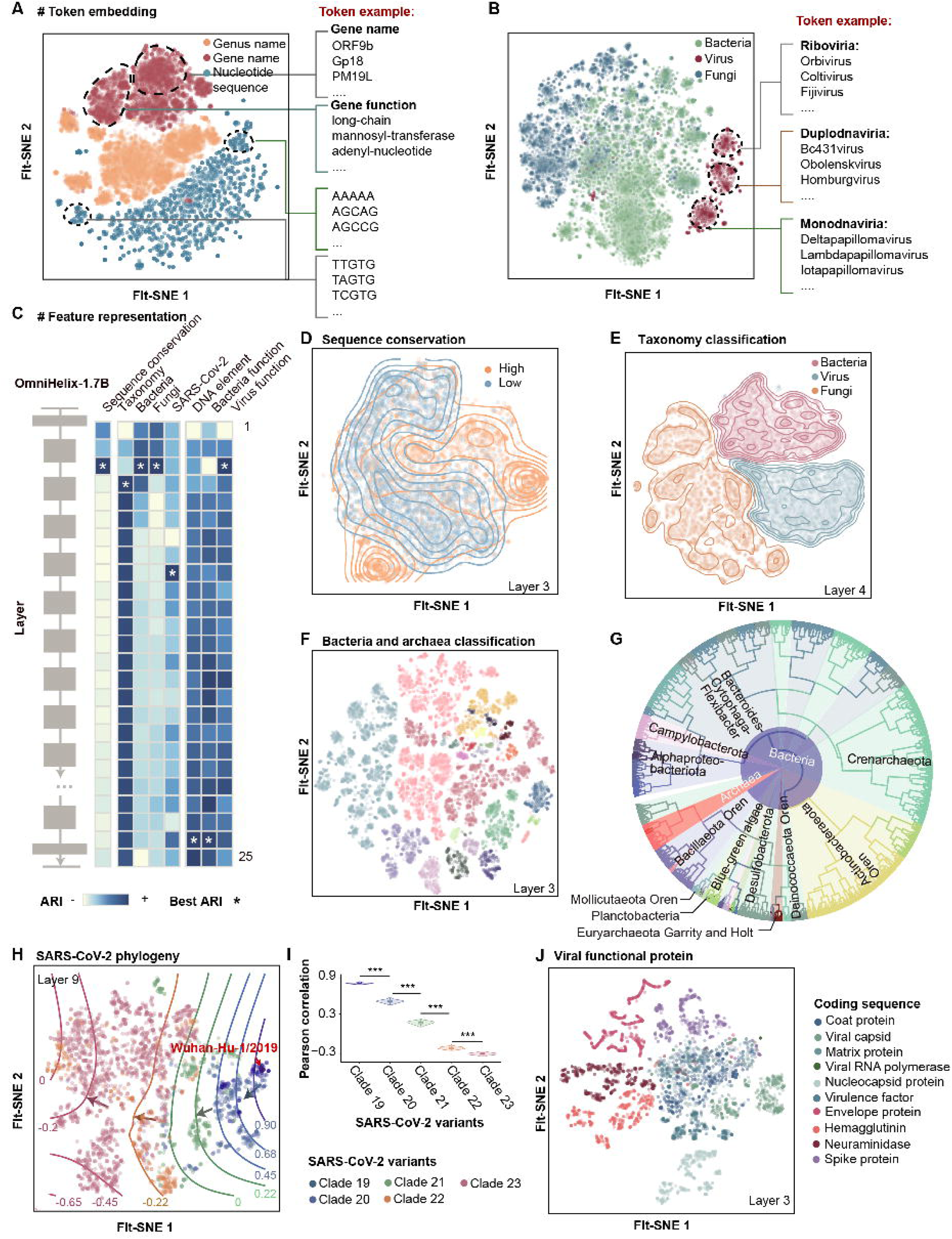
Sequence and token representations analysis. (**A-B**) t-SNE plots visualize the word embeddings of tokens with distinct semantics (**A**) and superkingdoms (**B**). Each point represents a word, color-coded by its semantic group (**A**) and superkingdom (**B**). (**C**) Heatmap illustrates the Adjusted Rand Index (ARI) values for transformer layers across different representation tasks. The ARI scores are scaled by task, with asterisks denoting layers exhibiting the highest AIR for each task. (**D**) t-SNE plot visualizes sequences with varying sequence conservation represented by transformer layer 2. Each point represents a sequence, color-coded by sequence conservation. The counter plot reflects the density distribution of sequences of distinct conservation. (**E**) t-SNE plot visualizes sequences from different superkingdoms represented by transformer layer 3. The counter plot depicting the density distribution of sequences from different superkingdom categorizations. (**F**) t-SNE plot visualizes bacterial sequences represented by transformer layer 2. The color-coded according to bacterial class. (**G**) Phylogenetic tree of the bacteria constructed with the similarity between sequence representations. Line segment colors denote bacterial class. The outer pie chart colors signify bacterial phylum, and inner pie chart colors distinguish between bacteria and archaea. (**H**) t-SNE plot visualizes sequences from different SARS-CoV-2 variants represented by transformer layer 8. Counter lines reflect the Pearson correlation of each strain with the first recognized strain (Wuhan-Hu-1/2019), with the red-highlighted point representing Wuhan-Hu-1/2019. (**I**) Violin and box plot illustrate the pearson correlation between different SARS-CoV-2 clades and the first recognized strain. Data is randomly sampled 1, 000 times from each branch, with 100 strains per sample, and the average Pearson correlation is shown. The boxplot presents the median, upper, and lower quartile of the distribution. (**J**) t-SNE plot visualizes virus sequences represented by transformer layer 2. \*\*\**p*-value < 0.001.

Our further exploration of the feature representation space extended to three specific tasks in terms of sequence conservation delineation, species differentiation and functional sequence differentiation. The ARI indexes underscore that different layers manifest optimal representation capabilities for distinct tasks (**Figure 3C**). Specifically, the second layer can segregate sequences with varying conservation levels (**Figure 3D**). The third layer can separate sequences from different superkingdoms (**Figure 3F**). Leveraging bacteria, archaea 16s RNA and fungi ITS sequences, the second layer enables species discrimination at class level (**Figure 3F** and **Supplementary** Figure 4). Phylogenetic trees constructed based on the sequence representations further confirm the model’s representation ability at both class and phylum levels (**Figure 3G** and **Supplementary** Figure 4). The eight layer distinguishes different branches of the evolutionary progress of SARS-CoV-2 (**Figure 3H**). The phylogenetic trees underscore the model’s ability to recover the evolutionary trajectory (**Supplementary** Figure 4). Quantitatively, the similarity with the first recognized strain (Wuhan-Hu-1/2019) significantly diminishes with the evolution of branches (**Figure 3I**). The third and twenty-third layers can separate coding sequences of different functional proteins of viruses and bacteria, respectively (**Figure 3J** and **Supplementary** Figure 4). t-SNE visualization demonstrates the twenty-third layer’s capability in separating genomic elements based on their distinct functional roles (**Supplementary** Figure 4). Collectively, these findings highlight OmniNA-1.7B’s capacity to assimilate biological knowledge, encompassing both textual patterns and nucleotide sequences.

## Comprehensive evaluation of OmniNA’s generalization across **nucleotide sequence tasks**

To assess the generalization capabilities of OmniNA on nucleotide sequence tasks, we conducted fine-tuning experiments on 17 diverse tasks encompassing genome and species detection. OmniNA generally achieved better F1 scores than benchmark methods (**Figure 4**). Specifically, for tasks related to human DNA element identification, OmniNA achieved comparable or superior performance as compared with the benchmark methods (**Figure 4A**). Besides, OmniNA-1.7B excelled in PAS, TIS signal, and splice site detection, achieving remarkable F1 scores of 0.97, 0.88, and 0.98, respectively (**Figure 4B-C**). The superior performance of the OmniNA-1.7B model can also be observed for gRNA off-target detection with an F1 score of 0.98 (**Figure 4D; Supplementary** Figure 5). In methylated region detection tasks, the OmniNA-1.7B model achieved comparable F1 and higher precision scores across various cell types compared to the 66M and 330M models (**Figure 4E**). Additionally, for chromatin features prediction, OmniNA-1.7B delivered high-accuracy predictions for chromatin accessibility and histone modification, surpassing current state-of-the-art methods (**Figure 4F**, **Supplementary** Figure 5 **and Supplementary Table 5**). Our method further demonstrated high performance in the prediction for TF occupation, achieving a median F1 score of 0.73 across all TF targets (**Figure 4F**, **Supplementary** Figure 5 and **Supplementary Table 5**). Besides, the model’s performance on pathogenic variation detection tasks improved with increasing model size (**Figure 4G**). We compared the OmniNA-1.7B model against the EVE method for detecting pathogenic variations based on multiple sequence alignment (MSA) (**Figure 4H**). Across 17 genes, OmniNA-1.7B achieved higher F1 scores in 12 genes compared to EVE. OmniNA also exhibited proficiency in species detection tasks (**Figures 4I-K**). In the pathogenic virus detection task (**Figure 4I**), the F1 scores of OmniNA-1.7B surpassed those of benchmark methods. Furthermore, on bacteria taxonomy classification tasks related to class, family, order, and phylum (**Figures 4J-K** and **Supplementary** Figure 5), OmniNA-1.7B outperformed the benchmark model. All detailed quantitative results are listed in **Supplementary Table 6**.

**Figure 4.**
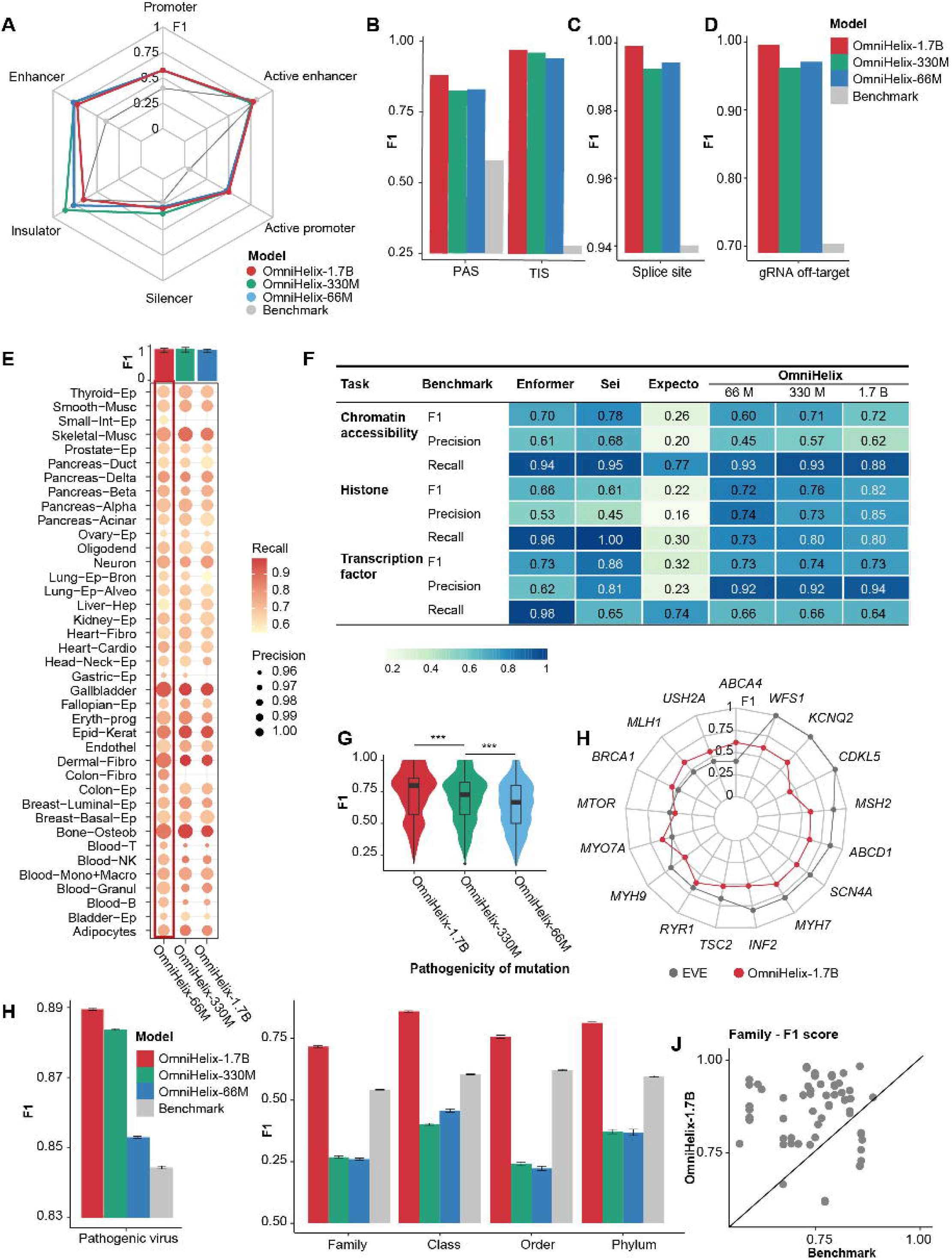
Performance of fine-tuned OmniNA on nucleotide sequence identification tasks. (**A**) Radar plot shows the F1 score achieved by OmniNA and the benchmark method on the task of DNA element detection. (**B-D**) Bar plot illustrating the F1 score achieved by OmniNA model and the benchmark method on the task of splice site detection (**B**), PAS/TIS site detection (**C**) and gRNA off-target prediction (**D**). (**E**) Dot plot depicts the performance of OmniNA with different parameters on the task of methylated region detection. The circle size reflects the precision score of method, and the color reflects the recall of the method. The bar plot in the above panel shows the F1 score of the OmniNA model with different parameters. (**F**) The performance of OmniNA and three benchmark methods, i.e. Enformer, Sei and Expecto on chromatin feature detection tasks. (**G**) The boxplot presents the median, upper, and lower quartile of the f1 score for the pathogenic variation detection performance across all genes. (**H**) Radar plot shows the F1 score of the OmniNA-1.7B model and benchmark method for pathogenic SNV detection. (**I**) Bar plot illustrating the F1 score achieved by OmniNA and the benchmark method on the task of pathogenic virus detection. Data is randomly sampled 1, 000 times from each branch, with 1, 000 strains per sample, and the average Pearson correlation is shown. The center line represents the mean, and the error bar indicates the 95% CI. (**J**) Bar plot illustrating the F1 score achieved by OmniNA and the benchmark method on the task of bacteria species classification. Data is randomly sampled 1, 000 times from each branch, with 1, 000 strains per sample, and the average Pearson correlation is shown. The center line represents the mean, and the error bar indicates the 95% CI. (**K**) The F1 score comparison between the OmniNA-1.7B model and benchmark method on bacteria phylum classification task. \*\*\**p*-value < 0.001.

### Discriminating mutation effects in distinct genomic contexts with OmniNA

After fine-tuning OmniNA-1.7B on diverse nucleotide learning tasks aligned with natural language paradigms, we further verify the model’s ability to predict the mutation impact within varied genomic contexts. We defined the effect score of a mutation as the OmniNA-1.7B estimated LL between mutated and wild-type nucleotide sequences (**Figure 5A**). Our initial focus was on mutations influencing PAS and TIS signals. Analyzing mutations in genomic loci around PAS regions reveals a consistent increase in the odds ratio (OR) between pathogenic and benign mutations as predicted mutation effect increasing (**Figure 5B**, Pearson correlation = 0.92, *p*-value = 9e-3). Similarly, mutations at TIS-associated genomic loci exhibit a proportional rise in the OR between pathogenic and benign mutations with escalating predicted mutation effects (**Figure 5C**, Pearson correlation = 0.92, *p*-value = 1e-2). Furthermore, OmniNA-1.7B captures the mutation effects at the 23 positions of CRISPR-Cas13 gRNA, showcasing significant alignment with actual mutation effects (Pearson correlation = 0.65, *p*-value = 6.5-4). Moreover, OmniNA-1.7B showcases discriminative power in identifying pathogenic effects at functional regions within protein coding sequences. For instance, the mutation effect in the domain region of the *BRCA1* coding sequence significantly surpasses than that in non-domain regions (**Figures 5D-E**; *p*-value < 1e-5). Structurally, the pathogenic effects across three domains outperform those in surrounding ordered and disordered structures (**Figure 5F**). Furthermore, we extend this discriminative capability to *CHD7*. The mutation effect in the domain region of CHD7 exhibits a significant elevation compared to non-domain regions (**Figures 5G-H**; *p*-value < 1e-5). This trend is consistent with the structural analysis, where the pathogenic effects across the three domains of CHD7 surpass those found in surrounding regions (**Figure 5I**). In conclusion, OmniNA-1.7B exhibits a grasp of the structural and functional intricacies of nucleotide sequences. This enables precise predictions regarding how mutations alter fundamental aspects, showcasing its potential in mutation effect prediction.

**Figure 5.**
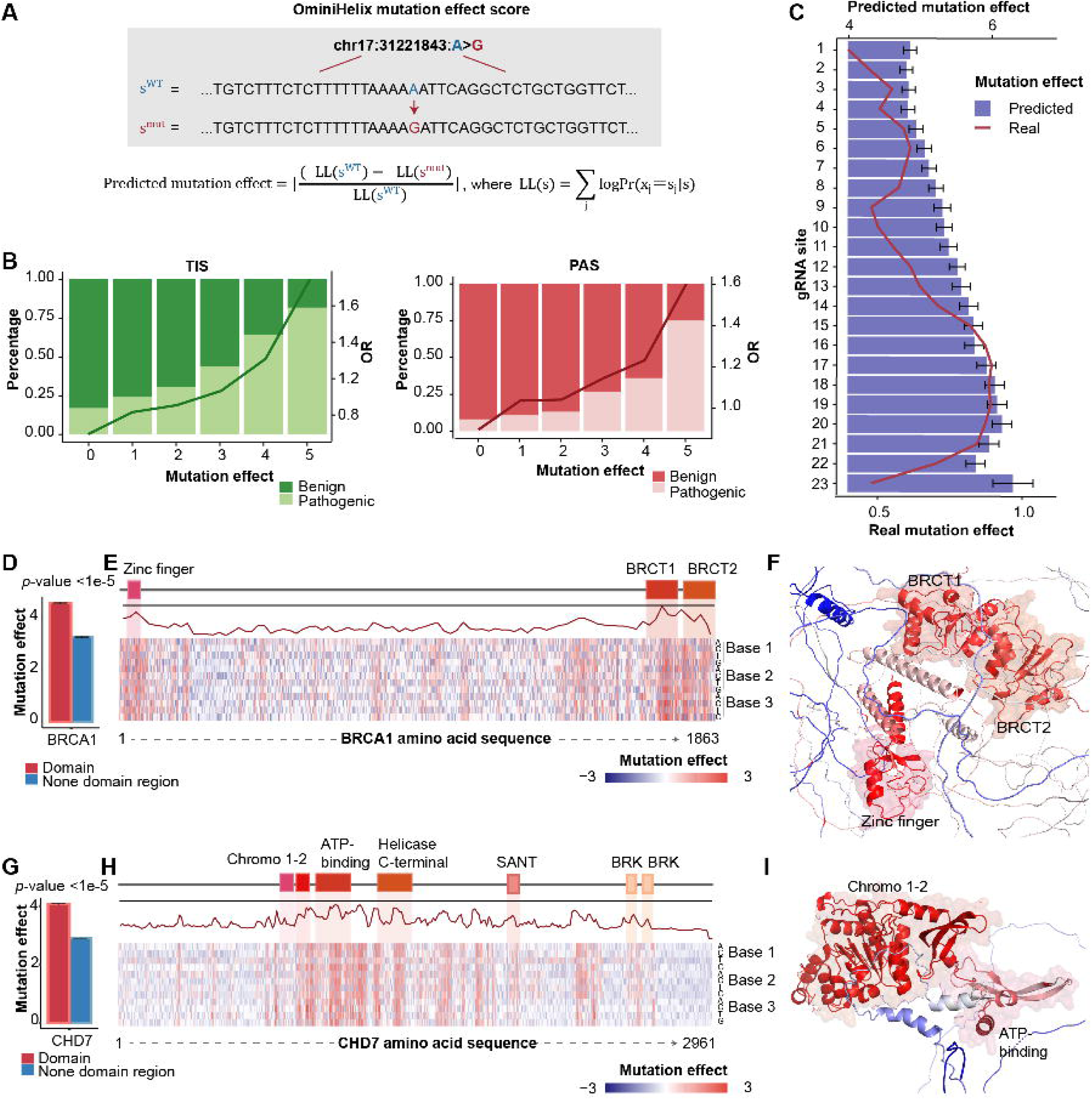
Predicted mutation effects of clinically and functionally relevant missense variants. (**A**) Schematic diagram of the mutation effect score calculation. (**B**) Bar plot illustrates the correlation between mutation effect scores and the proportion of pathogenic and non-pathogenic mutations. Line plot depicts the odds ratio (OR) between pathogenic and benign mutations. (**C**) Bar plot shows the mean predicted mutation effects for each gRNA position, with error bar representing the 95% confidence interval (CI) of the mutation score. (**D**) Bar plot shows the mean mutation effect score for mutations in the domain region compared to the non-domain region of *BRCA1* coding gene. The error bar represent the 95% CI of the distributions of mutation score. (**E**) Upper panel, schematic representation of the domains and non-domain protein coding regions of *BRCA1*; middle panel, maximum mutation effect score observed for each nucleotide base; lower panel, heatmap illustrating mutation effect scores across all possible 3 × 4 missense variants. (**F**) AlphaFold prediction of the three-dimensional structure of the BRCA1 protein. The residues are colored by the predicted mutation effect scores. (**G**) Bar plot shows the mean mutation effect score for mutations in the domain region compared to the non-domain region of *CHD7* coding gene. The error bar represents the 95% CI of the distributions of mutation score. (**H**) Upper panel, schematic representation of the domains and non-domain protein coding regions of *CHD7*; middle panel, maximum mutation effect score observed for each nucleotide base; lower panel, heatmap illustrating mutation effect scores across all possible 3 × 4 missense variants. (**I**) AlphaFold prediction of the three-dimensional structure of the CHD7 protein. The residues are colored by the predicted mutation effect scores.

### Ablation study for few-shot learning and cross-species generalization analysis

We conducted an extensive evaluation of the OmniNA’s few-shot learning capabilities. The larger model consistently outperformed its counterparts as the percentage of training data increased (**Supplementary** Figure 6). Notably, the OmniNA-1.7B model demonstrated comparable performance with 75% and 100% of the data, showcasing its effectiveness in learning histone occupation profiles from limited information. To further assess OmniNA’s cross-species generalization capability, we fine-tuned OmniNA on human DNA element classification tasks and applied it for mice DNA elements classification. As illustrated in **Supplementary** Figure 6, OmniNA achieved comparable performance across all evaluations with the benchmark methods trained specifically with mice sequences. Quantitative results are detailed in **Supplementary Table 7**. These findings underscore OmniNA’s robust few-shot learning and impressive generalization capacities.

## Discussion

Deep learning is undergoing a paradigm shift to the foundation model, which can generalize to a wide range of downstream tasks and result in emergent capabilities. In the realm of nucleotide sequencing, the understanding of the intricate rules governing sequences is paramount, and aligning various nucleotide learning tasks with natural language is an emerging need. In this context, the creation of our foundation model OmniNA not only responds to the technical exigencies posed by the data deluge but also addresses the inherent limitations of traditional bioinformatic methodologies. By deploying a foundation generative model, our objective is to empower biologists and researchers with a versatile tool capable of autonomously discerning intricate relationships within nucleic acid sequences. This model not only facilitates the effective capture of inter-sequence correlations but also provides a robust foundation for subsequent downstream tasks.

We observed a positive correlation between the model size and OmniNA’s generation capabilities. As model size increased, we noted a corresponding increase in the fidelity of generated annotations. The diversity of the generations is also increased as the model size enlarged, particularly in generations with low-frequency words encountered during the pre-training process. The expansive capacity of larger models to effectively handle a broader range of vocabulary and learn from low-frequency words contributes to their superior generative performance. Our exploration extended to the realm of reproducibility across different truncations of the same sequence. We proved that various truncations of a single sequence consistently produced similar representations, emphasizing the stability of the model’s annotation generation. Furthermore, the generated annotations exhibited a degree of overlap for genomes with overlapping regions, highlighting the model’s capacity to capture the intricacies of shared genomic elements. In summary, our findings underscore the advantages of larger models, not only in terms of fidelity but also in the diversity and stability of generations.

Our study presented evidence for that OmniNA learns representations of tokens and sequences across diverse biological contexts. In the absence of supervision, our model exhibits a capacity to reveal nuanced sensitivity to different biological structures and evolutionary information. As transformer layers deepen, the model’s aptitude to represents complex biological information becomes more evident. Shallow layers appear proficient in capturing elementary information, such as sequence conservation and species distinctions. The intermediate layers demonstrate a heightened capacity to discern the evolutionary dynamics of the SARS-Cov-2. The deeper layers capture information pertaining to sequence functionality. Our study represents a grasp of understanding the unsupervised acquisition of biological knowledge by LLMs.

Unlike canonical deep learning approaches in genomics that learn from a specific task, OmniNA is a general solution that can be applied to a wide range of tasks across species classification, DNA and RNA processing. This versatility of handling a varying number of tasks without the need for task-specific architectures reduces the complexity and effort required in designing and training specialized models. The flexibility of large-language models in accommodating various tasks in a QA manner enables a broader range of users, including those with less specialized expertise in deep learning, to harness the power of advanced language models for diverse applications ^40^. In our evaluation of the fine-tuning OmniNA on 17 tasks, the model demonstrated comparable or superior performance to dedicated deep learning models in specific tasks, highlighting its adaptability in genomic contexts. Importantly, our findings suggest that, while different-scale models achieve similar results in straightforward tasks such as DNA element detection, the superior performance of larger-scale models becomes evident in more challenging tasks like pathogenic mutation identification. This scalability feature positions OmniNA as a formidable tool, demonstrating its ability to learn intricate nucleotide sequence grammar and translate it into mutation effect prediction. This ability enhances its utility in understanding the functional implications of genetic variations. In summary, OmniNA emerges as a versatile and scalable solution, empowering users across various expertise levels to navigate genomics with ease and efficacy.

This study has several limitations. First, the scaling on the performance of deep learning models shows the existence of power laws between the model and data sizes and the performance of the system ^41^. As genomics research advances, incorporating OmniNA with larger model sizes into the evolving landscape of high-throughput sequencing technologies necessitates considerations and adaptations. Second, the model’s fine-tuning effectiveness may be sensitive to the quality and completeness of the genomic data. In instances where the labeled data are either incomplete or biased, OmniNA’s performance could encounter limitations. Addressing this challenge involves continuous efforts in refining genomic annotations to align with evolving biological knowledge. Third, in comparison with state-of-the-art methods, it is crucial to recognize potential overlaps between OmniNA’s test set and the training data of other models. Besides, for regression models, including MPRA-DragoNN, Enformer and Expecto, we employed optimal AUC values from training data to determine cutoffs for converting tasks into classification tasks. These factors could introduce evaluation biases. Lastly, OmniNA is currently limited to learning sequences of approximately 3000 bps. Sequences exceeding this length require truncation for effective learning. Future iterations of OmniNA will incorporate an extended receptive field to facilitate long-range interaction learning, ensuring its applicability to a broader spectrum of genomics tasks.

In conclusion, OmniNA represents an endeavor in leveraging foundation models for comprehensive nucleotide learning across diverse species and genome contexts. Its alignment with natural language, enhances our understanding of nucleotide sequence rules, and provides a versatile and user-friendly platform for addressing challenges in genomics and molecular biology. As we open-source for collaborative exploration, we anticipate that OmniNA will open avenues for uncovering universal principles governing nucleotide sequences, offering novel insights and methodologies for exploring the intricacies of genomics.

## Acknowledgment

This work was supported by the National Natural Science Foundation of China [32270688 and 31801117]; the Program for Changjiang Scholars and Innovative Research Team in University in China [IRT_14R40]; the Tianjin Science and Technology Committee Foundation [17JCYBJC25300]; and the Chinese National Key Research and Development Project [2018YFC1315600]. We want to thank all the researchers for their generosity to make their data publicly available.

## Author contributions

X.L. designed and supervised the study. X.L. and X.S. wrote the manuscript.

X.L. and X.S. revised the manuscript. X.L. and X.S. collected the data. X.S. processed the data. X.L. and X.S. interpreted the results. All authors reviewed and approved the submission of this manuscript.

## Supplementary Figure and Table legends

**Supplementary Figure 1. OmniNA model characteristics and training dynamics during pre-training.** (**A**) Depiction of OmniNA model details, including layer configurations, head numbers, and hidden sizes for various model sizes. (**B**) Illustration of the training loss trajectory throughout the pre-training procedure.

**Supplementary Figure 2. Schematic representation of OmniNA model architecture**.

**Supplementary Figure 3. Fidelity assessment of OmniNA’s generations. (A)** Bar plot illustrates the cosine similarity between generated and ground truth sequences across coding sequences of diverse proteins. The grey line denotes the sequence conservation for the proteins. The dashed line indicates the cutoff for sequences with high or low conservation. (**B**) Confusion matrix detailing results for species, genomic type, and enzyme datasets.

**Supplementary Figure 4. Visualization of sequence representations.** (**A**) t-SNE plot presenting virus sequences represented by transformer layer 2. The dots are color-coded by virus class. (**B**) Phylogenetic tree of viruses constructed based on sequence representation similarities, with line segment colors denoting virus class and pie chart colors signifying virus phylum. (**C**) Phylogenetic tree of SARS-CoV-2 constructed similarly, with line segment and pie chart colors indicating SARS-CoV-2 clade. (**D**) t-SNE plot showcasing the coding sequences of bacteria proteins represented by transformer layer 23. (**E**) t-SNE plot illustrating DNA elements sequences represented by transformer layer 23.

**Supplementary Figure 5. Detailed assessment of fine-tuned model performance across diverse targets.** (**A**) Radar plot depicting gRNA efficiency for each gene.(**B**) Bar plot displaying the F1 score for each chromatin accessibility target. (**C-D**) Confusion matrix representing histone occupation (**C**) and transcription factor binding dataset (**D**). (**E**) The F1 score comparison between the OmniNA-1.7B model and benchmark method on bacteria phylum classification task.

**Supplementary Figure 6. Analysis of few-shot learning and generalization.** (**A**) Line plot illustrating the validation set loss with increasing training data percent in the histone occupation task. (**B**) Bar plot showcasing F1 scores achieved by OmniNA and benchmark methods in the mice DNA element detection task.

**Supplementary Table 1.** Prompts guiding the organization of pre-training and fine-tuning data.

**Supplementary Table 2.** Datasets utilized for the fine-tuning process.

**Supplementary Table 3.** Data sources from the ENCODE database for the detection of chromatin-accessible regions, histone-occupied regions and transcription factor binding regions.

**Supplementary Table 4.** Compilation of benchmark methods employed for evaluation.

**Supplementary Table 5.** F1 scores corresponding to each target in chromatin accessibility, histone occupation, and transcription factor binding analyses.

**Supplementary Table 6.** Results from multi-task fine-tuning experiments.

**Supplementary Table 7.** Evaluation of cross-species generalization capability.

